# Synaptic status differentially regulates neurofilaments in dendritic spines

**DOI:** 10.1101/2022.09.07.506909

**Authors:** Clara-Marie Gürth, Maria Augusta do Rego Barros Fernandes Lima, Angel Rafael Cereceda Delgado, Victor Macarrón Palacios, Jasmine Hubrich, Elisa D’Este

**Affiliations:** Department of Optical Nanoscopy, Max-Planck-Institute for Medical Research, Heidelberg, Germany; Department of NanoBiophotonics, Max-Planck-Institute for Multidisciplinary Sciences, Göttingen, Germany; Optical Microscopy Facility, Max-Planck-Institute for Medical Research, Heidelberg, Germany

**Keywords:** Neurofilaments, synapses, dendritic spines, STED nanoscopy

## Abstract

Neurofilaments are one of the main cytoskeletal components in neurons and they can be found in the form of oligomers at pre- and postsynapses. How their presence is regulated at the postsynapse remains widely unclear. Here we systematically quantified by immunolabeling the occurrence of the neurofilament isoform triplet neurofilament light (NFL), medium (NFM), and heavy (NFH) at the postsynapse with STED nanoscopy together with markers of synaptic strength and activity. Our data shows that within dendritic spines neurofilament isoforms rarely colocalize with each other and that they are present to different extents, with NFL being the most abundant isoform. The amount of the three isoforms correlates with markers of postsynaptic strength and presynaptic activity to varying degrees: while NFL shows moderate correlation to both synaptic traits suggesting its involvement in synaptic response, NFM and NFH were only correlated on a low level. By quantifying the presence of neurofilaments at the postsynapse within the context of the synaptic status, this work sheds new light on the regulation of synaptic neurofilaments and their possible contribution to psychiatric disorders.

## 1. Introduction

Neurofilaments (NFs) are among the main components of the cytoskeleton in neurons. They are heteropolymers of NF-light (NFL), NF-medium (NFM), and NF-heavy (NFH), and can further incorporate into their backbone either α-internexin or peripherin in the central or peripheral nervous system, respectively [1]. NFM and NFH have long tails enriched in phosphorylation sites that protrude into the periphery of the filament [for reviews 2–4]. Variability in the subunit stoichiometry and the dynamic phosphorylation landscape indicate NFs have a high potential for being finely regulated [5]. Indeed, these features are differentially modulated based on the cell type, developmental stage, and subcellular compartment [2,3].

For a long time NFs have been considered as mere space-fillers in the axon, albeit present also in dendrites [6–8]. Only in the last years a link between NF function in synapses and psychiatric disorders started to emerge [9–12]. Indeed, NF were identified at pre- and postsynaptic sites of excitatory and inhibitory synapses where they have a different phosphorylation status compared to the overall NF population [13–15]. Interestingly, all subunits appear more concentrated in the postsynaptic compartment than at the presynapse. Inhibition of NFH was shown to depress long-term potentiation (LTP) in the hippocampus without altering the spines morphology, which, on the other hand, is affected in NFL knockout animals [13,16]. Furthermore, NFM functionally interacts with dopamine D1-receptors, while both NFL and α-internexin regulate the localization of N-methyl-D-aspartate (NMDA) receptors subunits, providing direct evidence for the involvement of NFs in the regulation of synaptic functions [12,13,16–19]. However, whether and how the NF composition of a spine correlates with the synaptic status, and in particular the presynaptic activity and the postsynaptic strength, is unclear. Answering this question would reveal whether NFs play a role in the short-term response to synaptic activity or rather in the structural rearrangements occurring during LTP.

To clarify the involvement of NFs in postsynaptic activities, we used quantitative multicolor super resolution stimulated emission depletion (STED) microscopy of the NF triplet NFL, NFM, or NFH in combination with proxies of either postsynaptic strength (homer) or of presynaptic activity (synaptotag-min-1, Syt1, live-cell uptake [20,21]) on cultured hippocampal neurons. We found that NFL is the most abundant isoform in dendritic spines. Overall the amounts of NF isoforms correlated with both homer and Syt1 signal to varying extent with NFL showing the highest and NFM the lowest correlation, suggesting that its occurrence is influenced by the postsynaptic strength and activity.

## 2. Materials and Methods

### 2.1 Preparation of neuronal cultures

All procedures were performed in accordance with the German Animal Welfare Act (Tierschutzgesetz der Bundesrepublik Deutschland, TierSchG) and the Animal Welfare Laboratory Animal Ordinate (Tierschutz-Versuchstierverordnung, TierSchVersV) according to which no specific ethical authorization or notification is required. The sacrificing of P0–P2 rats was supervised by animal welfare officers of the Max Planck Institute for Medical Research (MPImF) and conducted and documented according to the guidelines of the TierSchG (permit number assigned by the MPImF: MPI/T-35/18).

Primary hippocampal or cortical neuronal cultures were prepared from dissociated tissue of P0-P2 postnatal wild-type Wistar rats of either sex (Janvier-Labs) as described previously in [21]. Briefly, dissected hippocampal tissue was digested with 0.25% trypsin for 20 minutes at 37°C, dissociated, and maintained in Neurobasal supplemented with 2% B27, 1% GlutaMAX and 1% penicillin/streptomycin (all from Gibco, Thermo Fisher Scientific). Cells were seeded at a concentration of 110,000/well in 12-well plates on ø 18mm glass coverslips coated with 0.1 mg/ml poly-ornithine (Sigma-Aldrich) and 1 μg/ml laminin (Corning). Medium was changed to fresh supplemented Neurobasal 1-2 hours after seeding and cultures were maintained in an incubator (37°C, 5% CO_2_, 95% rH) until day *in vitro* (DIV) 15-25 without inhibition of glial cell growth by AraC.

### 2.2 Sample preparation and immunostaining

Cultured neurons were transduced at DIV 7 for volume labeling with AAV-hSyn-EGFP, which was a gift from Bryan Roth (Addgene #50465). The preparation of adeno-associated viral vectors was performed as previously described in [22] and purified as described in [23]. Turnover of synaptic vesicles in mature cultures (DIV 15-25) was determined by live labeling with Atto647N-labeled mouse antibody against the luminal domain of synaptotagmin-1 (Synaptic Systems, cat. 105 311AT1, 1:500 in culture medium) for 1 hour. Samples were afterwards washed three times in prewarmed ACSF (126 mM NaCl, 2.5 mM KCl, 2.5 mMCaCl_2_, 1.3 mM MgCl_2_, with 30 mM Glucose, 27 mM HEPES) before fixing.

Mature (DIV15-25) neuronal cultures were fixed in 4% PFA in PBS, pH 7.4, quenched with quenching buffer (PBS, 100 mM glycine, 100 mM ammonium chloride), permeabilized for 5 min in 0.1% Triton X-100 in PBS and blocked with 1% BSA in PBS for 1 hour. Samples were then incubated for 1 hour at room temperature with primary antibodies, secondary antibodies and phalloidin in PBS, each followed by five washes in PBS. Primary antibodies used were: homer1 guinea pig (Synaptic Systems, cat. 160 004, 1:500 dilution), neurofilament-L rabbit (Synaptic Systems, cat. 171 002, 1:200 dilution), neurofilament-L chicken (Synaptic Systems, cat.171 006, 1:200 dilution), neurofilament-M rabbit (Synaptic Systems, cat. 171 203, 1:200 dilution), neurofilament-M mouse (Synaptic Systems, cat. 171 211, 1:200 dilution), neurofilament-H rabbit (Synaptic Systems, cat. 171 102, 1:200 dilution), neurofilament-H mouse (Synaptic Systems, cat. 171 111, 1:200 dilution). Note that antibodies against the same neurofilament isoform were raised against the same epitope of the proteins. Secondary antibodies, nanobodies, and dyes used were: Star635P anti-guinea pig (Abberior, cat. 2-0112-007-1, 1:100 dilution) and Alexa Fluor 594 anti-rabbit (Thermo Fisher, cat. A-21207, 1:100 dilution), Alexa Fluor 488 anti-chicken (Thermo Fisher, cat. A-21467, 1:100 dilution), Star635P anti-mouse (IgG2) (NanoTag, cat. N2702-Ab635P-S, 1:200 dilution), Star580 anti-mouse (IgG1) (NanoTag, cat. N2002-Ab580-S, 1:200 dilution), phalloidin Alexa Fluor 405 (Thermo Fisher, cat. A30104; 1:200 dilution). Samples were afterwards embedded in Mowiol® supplemented with DABCO and cured for at least 1 hour at room temperature before imaging.

### 2.3 Confocal and STED Imaging

Samples were imaged on an Abberior Expert Line Microscope (Abberior Instruments GmbH) on a motorized inverted microscope IX83 (Olympus) and equipped with pulsed STED lines at 775 nm and 595 nm, excitation lasers at 355 nm, 405 nm, 485 nm, 561 nm, and 640 nm, and spectral detection. Spectral detection was performed with avalanche photodiodes (APD) and detection windows were set to 650-725 nm, 600-630 nm, 505-540 nm, and 420-475 nm to detect Atto647N/Star635P, Alexa Fluor 594/Star580, Alexa Fluor 488/EGFP and Alexa Fluor 405, respectively. Images were acquired either with a 20x/0.4 NA oil immersion lens with pixel size of 100 nm or with a 100x/1.4 NA oil immersion lens, 30 nm pixel size, and pinhole to 100 μm (1 A.U.). Laser powers and dwell times were kept consistent during the entire experiments.

### 2.4 Image processing and analysis

Images shown in figures were visualized and processed with Imspector (Abberior Instruments GmbH), FIJI (*https://fiji.sc* version 1.53f51, [24]) and MATLAB 2021b (MathWorks). Images are shown as smoothed data with a low pass Gaussian filter and 1-5% background subtraction. Brightness was adjusted uniformly throughout the images. Manual segmentation of spines was performed in FIJI, based on either the EGFP volume labeling (for single NF analyses) or on phalloidin labeling (for colocalization analysis). Spines crossed by bright axonal NF signal were excluded from the analysis. In addition to the spine segmentation, a second mask was generated for each raw image by applying an automatic Otsu’s threshold to STED images of synaptic markers or a 4-counts background threshold for STED images of NFs. Each image was then masked with the intersection between the defined spine regions and its respective second mask. Afterwards, for each region of interest (ROI), the area and the intensity of the enclosing pixels were computed. The amount was defined as the area times the mean intensity.

For comparing amounts of NF and synaptic markers, we used the standardized data sorted by experimental round. The standardization was computed as 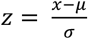, where *x* is the input data point, *μ* is the group mean and *σ* the group standard deviation. The Spearman’s correlation coefficient was then calculated for each NF isoform versus the synaptic marker, after grouping all the corresponding standardized experimental rounds. Spearman’s correlation coefficients were interpreted as suggested in [25].

Colocalization was performed using a custom code written in MATLAB. For analysis on the cellular level, NF signal of entire field of view was segmented from the background with a binary mask. The mask was generated after saturating 90% of the original confocal image, followed by an 8-pixel median filtering, to remove noisy pixels, and by a global Otsu’s threshold. On the spine level, we used the masks generated from the manual segmentation of STED images described above. Thereafter, the respective masks were applied to the original images and the Pearson’s correlation coefficient for non-zero pixels was computed for each pair of NF isoforms to characterize their degree of colocalization.

## 3. Results

### 3.1 NF isoforms poorly colocalize at the postsynapse

To analyze NFs at the postsynapse while preserving the isoform specificity and spatial resolution, our method of choice was immunofluorescence labeling. Therefore, we selected isoform-specific antibodies and firstly characterized the cellular distribution of the NF isoforms NFL, NFM, and NFH in hippocampal neurons using confocal microscopy. All three NF isoforms strongly labeled cultured neurons and exhibited a stronger signal in axons than in dendrites (**Figure 1A and S1A**). Interestingly, marked differences in the expression profile of each of the different isoforms were observed at cellular level, and in the distal axons in particular (**Figure S1A and S1B**). At higher magnification, NFM and NFH signals showed more characteristic filamentous organization within dendrites than NFL, whose signal was more granulated and dispersed throughout dendritic regions (**Figure S1C**). The differences in the expression pattern are confirmed by colocalization analyses: signals from the three different NF isoforms colocalized to only approximately 20-30% compared with the control experiments, in which the same isoform (NFM) was labeled with multiple fluorophores to compensate for chromatic artefacts (75%) (**Figure 1B and S1D**). Together with western blot analysis (**Figure S2**), this data confirms that the chosen antibodies are capable of detecting different isoforms and protein distribution, and suggests that different neuronal regions possess different NF expression patterns.

**Figure 1.**
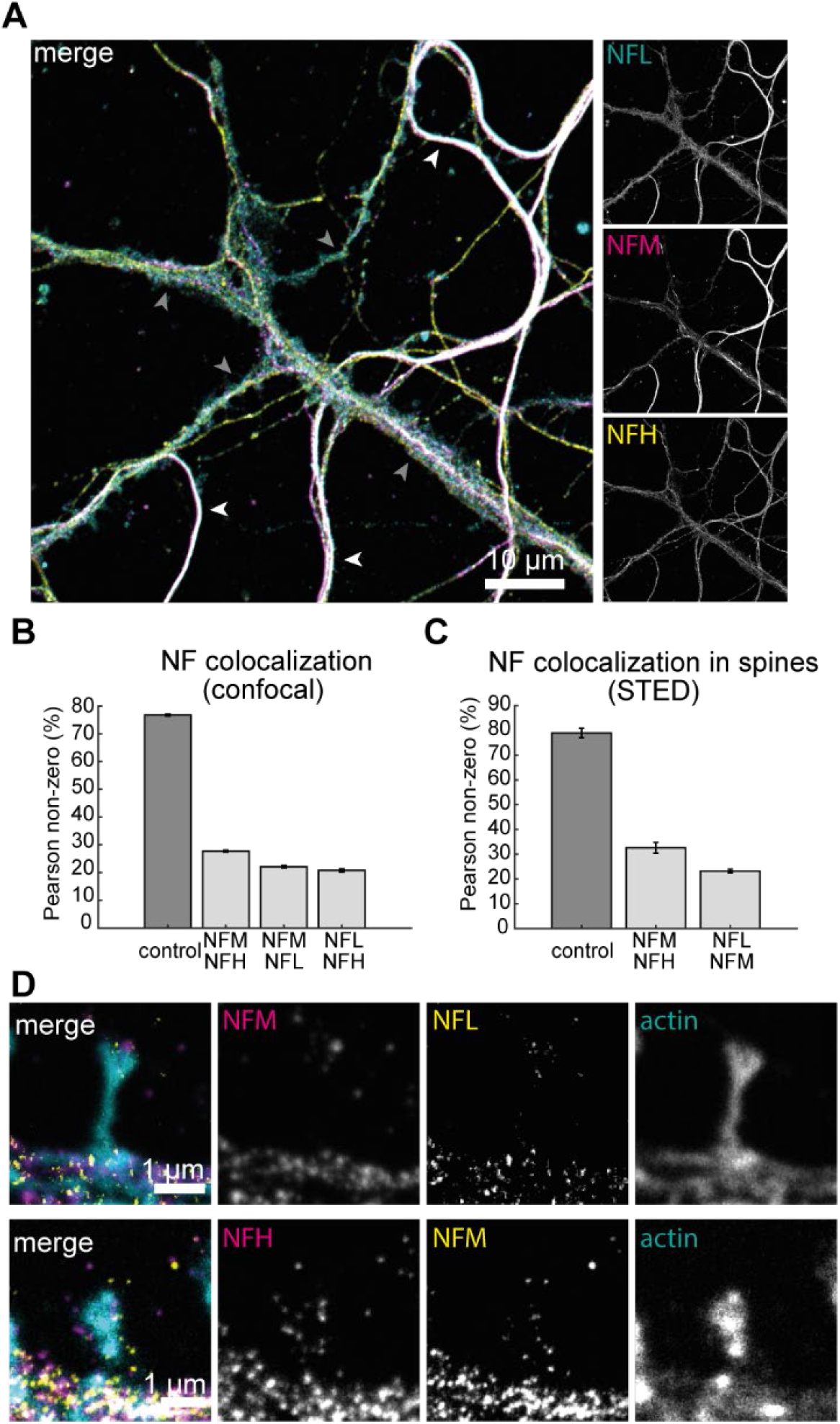
NF isoforms in hippocampal neurons. (A) Confocal images of the different NF isoforms NFL (cyan), NFM (magenta) and NFH (yellow) in primary hippocampal neurons. White arrowheads indicate axonal NF signal and grey arrowheads indicate dendritic NF signal. Scale bar is 10 μm. (B) colocalization analysis of NFL, NFM, and NFH on cellular level in confocal images as shown in (A) compared to control images with same isoform (NFM) labeled in all fluorescent channels. Shown is the mean values of Pearson’s correlation coefficient of all pixels with non-zero intensities. Error bars show the standard deviation of the mean (SEM). Data calculated from 21 different field of views. (C) Colocalization analysis of NFL, NFM, and NFH on the synaptic level in STED images as shown in (D) compared to control images with same isoform (NFM) labeled in both STED channels. Shown is the mean of the Pearson’s correlation coefficient of all pixels with non-zero intensities. Error bars show SEM. Data calculated from a total of 534 manually segmented spines from 3-5 different cells per condition. (D) Representative images of dendritic spines labeled with two NF isoforms (magenta/yellow) and actin (cyan). Scale bar is 1 μm.

Next, we focused on dendritic spines, to assess the role of NF isoforms at the postsynapse. To this aim, we performed dual-color STED imaging in neuronal cultures together with phalloidin, to visualize the spine morphology. Distinct, non-colocalizing puncta were observed in the spines when co-labeling NFL and NFM or NFH (**Figure 1D**). We further quantified this evidence by performing a colocalization analysis specifically at manually segmented dendritic spines in dual-color STED images (**Figure 1C**). We observed only a low degree (20-30%) of colocalization at the postsynapse for both NFL-NFM and NFM-NFH compared to the control (75%) with only one isoform (NFM) labeled with both fluor-ophores. This shows the sensitivity and specificity of our model system and that, although signal of all NF isoforms is present at most postsynapses, this signal rarely colocalizes.

### 3.2 NFL is the most represented isoform in dendritic spines

To further characterize the presence of NFs at the postsynapse, we next performed comparative STED imaging of either NFL, NFM, or NFH. To minimize the variability of the experiments, we chose primary antibodies raised in rabbit for all isoforms, so that the same secondary antibody could be used for all experiments. The epitopes recognized by these antibodies, which were validated by western blot (**Figure S2**), are the same as the antibodies used for the colocalization experiments shown in **Figure 1**. Both sets of reagents displayed similar labeling patterns of filamentous structures in the axons and puncta in dendritic spines (**Figure 2A**).

**Figure 2.**
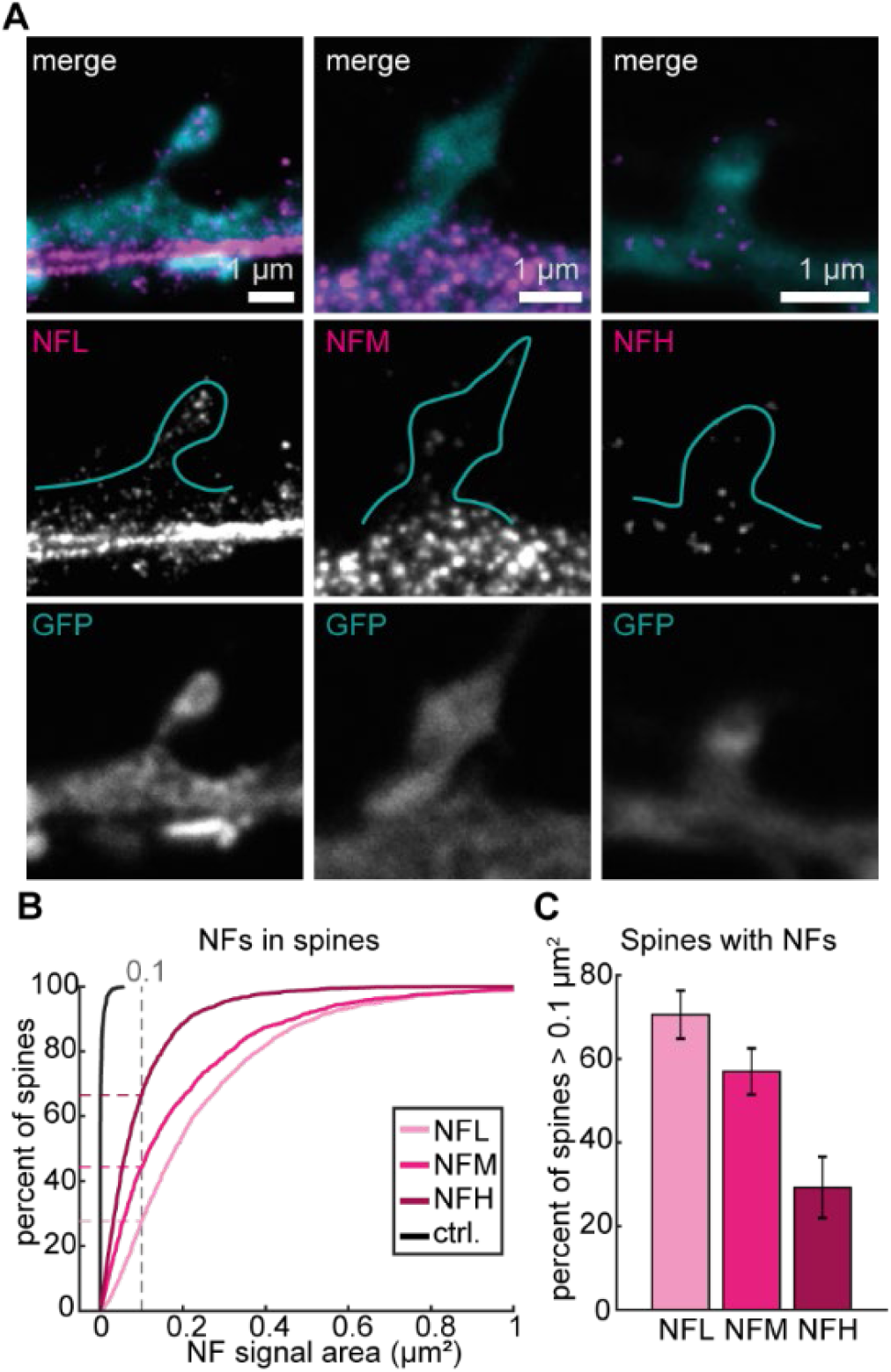
NF isoforms can be found at the postsynapse. (A) Representative STED images of dendritic spines with NFs (magenta) and volume labeling (confocal, cyan) with NFL (left), NFM (middle), or NFH (right). Spine shape as seen in volume labeling is represented in blue outline. Scale bars are 1 μm. (B) Cumulative distribution of spines containing NFs compared to control without primary antibody labeling. Dashed lines indicate individual values at a 0.1 μm^2^ threshold. (C) Percentage of spines containing more than 0.1 μm^2^ of NF signal. Data calculated from total 7660 spines (NFL = 2360, NFM = 1775, NFH = 2768, ctrl = 757) manually segmented from 9 independent experiments.

Next the presence of NFs inside dendritic spines was quantified. For this the intensity and area of NF signal inside the manually segmented spines, as identified by GFP volume labeling was analyzed (**Figure 2B**). The cumulative distribution of NF area within individual spines showed that in most of the spines, the area covered by NFL signal is larger than the area covered by NFM and NFH. Control images with only secondary antibody labeling showed low background signal compared to NF signal (**Figure 1B and S3**). Based on the cumulative distribution we tested the presence of NF signal area with a threshold of 0.1 μm^2^. We observed that around 70% of spines contained more than 0.1 μm^2^ NFL signal per spine whereas this value dropped to 55% and around 30% for NFM and NFH, respectively (**Figure 2C**). This demonstrates that NF isoforms are present at the postsynapse to varying extents and suggests a distinct role of NFL and NFM or NFH within dendritic spines.

### 3.3 NF isoforms in spines correlate to different extents with postsynaptic strength marker homer

After having quantified the NF content at the postsynapse we questioned whether the presence of NFs is in dependence of different synaptic traits. Firstly, we tested the connection with postsynaptic strength as also described previously [21]. We therefore additionally quantified the STED signal of the postsynaptic marker homer together with the STED signal of either NFL, NFM or NFH in manually segmented spines (Figure 3A). The amount of NF and the amount of homer were determined based on the area and mean intensity of the respective STED signal.

**Figure 3.**
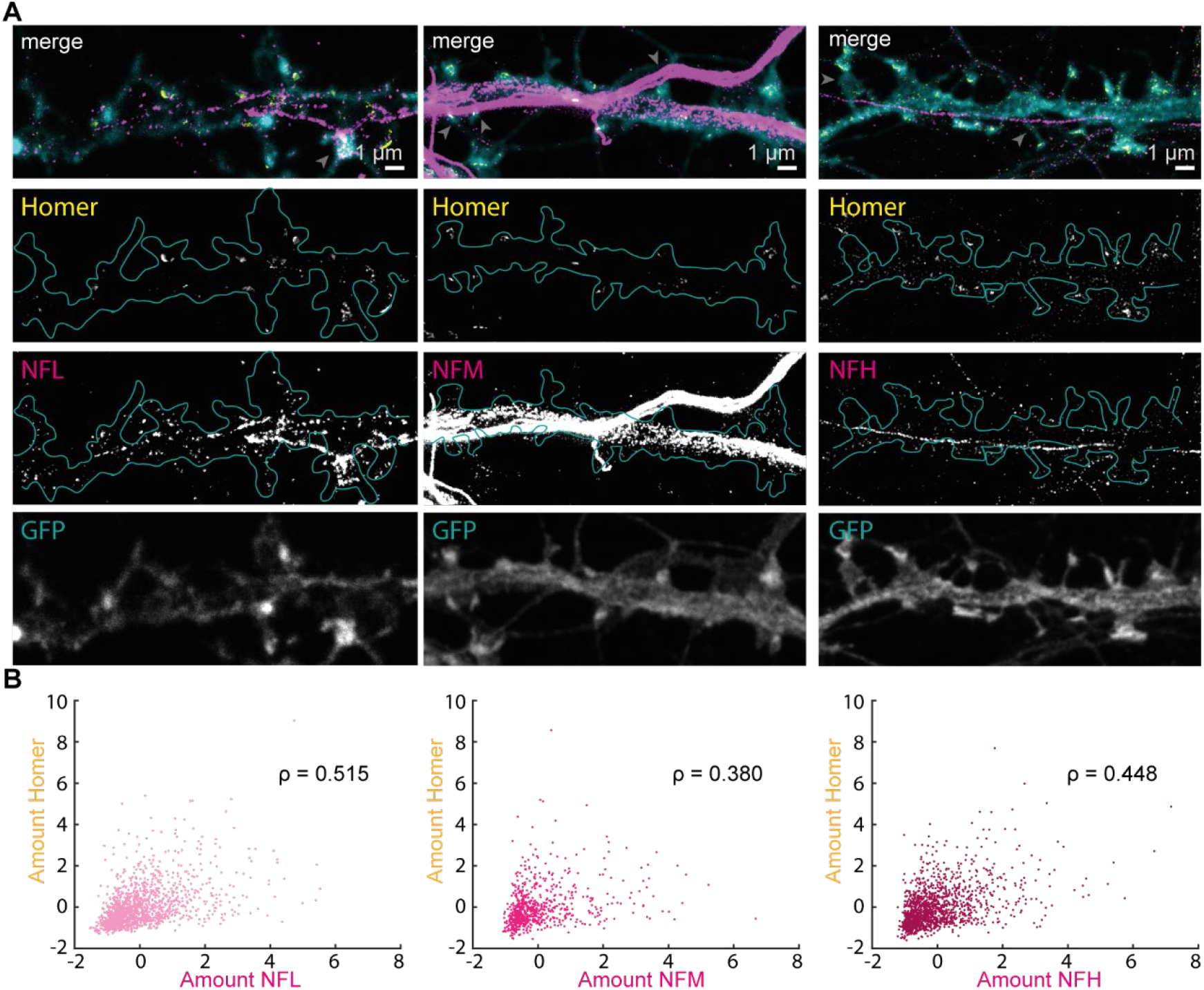
NFs at the postsynapse are differentially regulated by postsynaptic strength. (A) Representative dual-color STED images of dendrites with homer (yellow), NFs (magenta), and volume labeling (confocal, cyan) with NFL (left), NFM (middle), or NFH (right). Blue outline represents spine shapes on the dendrite as determined by volume labeling. Scale bars are 1 μm. Only signal above the 4-counts threshold used for data analysis is shown. Gray arrows in merge images point at examples of spines that were not considered for the analysis due to the overlay of strong axonal signal. (B) Scatterplot of NF versus homer protein amount (*area × mean intensity*). ρ indicates Spearman’s correlation coefficient with p values for NFL: 1.48e-101; NFM: 2.16e-27 and NFH: 2.93e-69. Data was obtained from 4637 manually segmented spines from 6 independent experiments, standardized to the mean of each experiment. Amounts represent arbitrary units.

The correlation between these two relative protein amounts exhibited moderate correlation between postsynaptic size and amount of NFL (ρ = 0.515) and only low correlation for NFM (ρ = 0.380) and NFH (ρ = 0.448) (**Figure 3B**). The presence of NFs is therefore differentially depending on postsynaptic strength as characterized by homer labeling with NFL having the strongest and NFM having the lowest correlation.

### 3.4 The amount of NFL at the postsynapse correlates with presynaptic vesicle recycling

Since the presence of NFs appeared to be influenced by postsynaptic strength to a differing degree depending on the isoform, we next questioned whether the presence of NFs in dendritic spines is further regulated by presynaptic activity. We therefore performed activity dependent live-labeling of Syt1, a proxy of synaptic activity, as previously described [20,21]. Specifically, immunolabeling of Syt1 was combined with immunolabeling of the three different NF isoforms as in previous experiments (**Figure 4A**). We then again correlated the amount of NFL, NFM or NFH with the number of actively recycled synaptic vesicles at individual synapses. We observed a moderate correlation between synaptic activity and the presence of NFL (ρ = 0.641), showing that more active synapses have larger amounts of NFL at their postsynaptic site (**Figure 4B**). Interestingly this correlation was lower for NFM and NFH amounts (ρ = 0.305 and 0.319, respectively). Hence NFL, the NF isoform that is most abundant at the postsynapse, shows a dependence not only to synaptic strength but also to synaptic activity.

**Figure 4.**
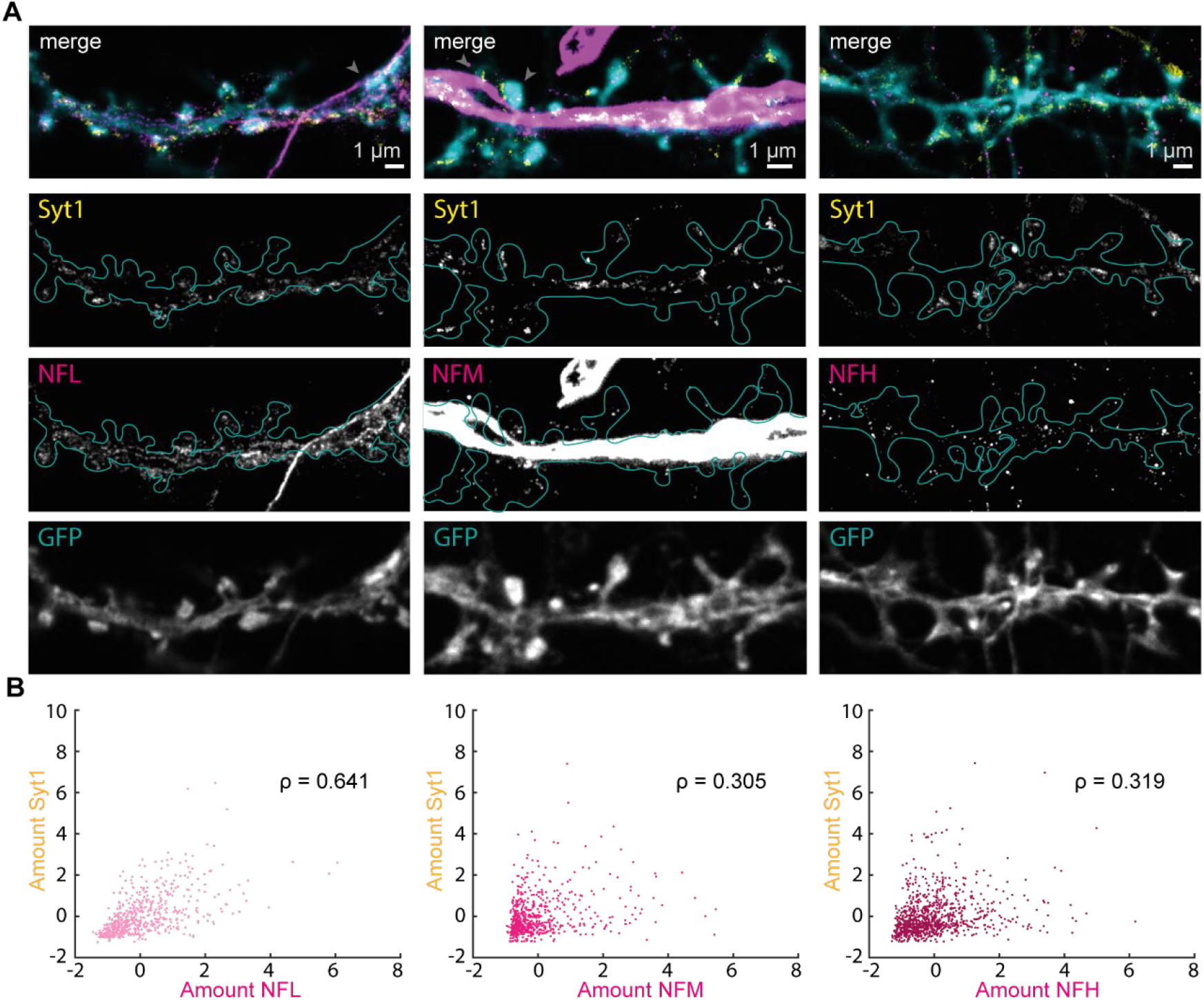
NFL at the postsynapse correlates with synaptic activity. (A) Representative dual-color STED images of dendrites with Syt1 (yellow), NFs (magenta), and volume labeling (confocal, cyan) with NFL (left), NFM (middle), or NFH (right). Blue outline represents spine shapes on the dendrite as determined by volume labeling. Scale bars are 1 μm. Only signal above the 4-counts threshold used for data analysis is shown. Gray arrows in merge images point at examples of spines that were not considered for the analysis due to the overlay of strong axonal signal. (B) Correlation scatterplot of NF protein amount and amount of labeled Syt1 (*area × mean intensity*). ρ indicates Spearman’s correlation coefficient with p values for NFL: 2.55e^-75^; NFM: 2.23e^-15^, and NFH: 1.26e^-23^. Data was obtained from 2657 manually segmented spines from 3 independent experiments, standardized to the mean of each experiment. Amounts represent arbitrary units.

## 4. Discussion

NFs belong to the most abundant proteins in neurons and are found in both axonal and dendritic regions. Although several studies suggest other NF functions beyond axonal structural support and connections with psychiatric disorders are known, their role at the synapse is only starting to emerge [9–11]. In the last decade, the presence of NF oligomers has been reported at postsynaptic sites, where they influence receptor function, LTP induction, and spine morphology [3,9]. In this work, we used quantitative multicolor nanoscopy of the NF triplet to characterize its distribution, and identify correlations between its presence and synaptic traits. In particular, we analyzed the correlation between NFL, NFM, and NFH with either a marker of postsynaptic strength (homer) or a marker of presynaptic activity (Syt1) [20,21].

The use of STED nanoscopy in combination with volume labeling enabled us to differentiate with high lateral resolution the postsynaptic sites from dendritic or axonal shafts. However, the possibility exists that signal coming from overlapping pre-axonal terminals is detected within the same focal volume, deteriorating the specificity of the results. Importantly, the abundance of NF is higher at the postsynapse than at the presynapse [3]. Furthermore, spines overlapping with obvious bright axonal signal were excluded from the analysis. Therefore, the impact of this technical limitation is minimized in this study.

Our study is based on chemical fixation and antibody labeling. Hence, the results are strongly dependent on the preservation of the structures, the specificity of the used reagents, and their ability to bind to the target structure. The influence of these aspects is commonly validated by live-cell imaging, which is however extremely challenging in the case of NFs due to the difficulties in tagging them while preserving their function [6]. Nonetheless, the axonal signal resembles the pattern observed by light and electron microscopy, and the punctate signal visualized by STED at postsynapses is in agreement with electron microscopy data showing the presence of oligomers at synaptic sites [6,13]. The capability of the antibodies to bind their target structure is relevant in the case of NFs, since they can be highly phosphorylated and the phosphorylation status of NF at synapses is distinctive compared to the rest of the neuron [9,26]. Hence, the affinity of antibody labeling might differ based on the post-translational modification and the possibility exists that not all NF molecules are being detected. While this might have an impact on absolute quantification of synaptic NFs this should only minorly affect the comparative correlation analyses as applied here.

Analysis of postsynaptic compartments revealed NF signal in the majority of dendritic spines. In these compartments, the area occupied by the fluorescent signal is larger for NFL than NFM and NFH. This difference cannot be ascribed to different optical resolutions, since the same fluorophores and imaging conditions were utilized, and therefore represent a broader distribution of the antibody labeling. The higher abundance of NFL in dendritic spines supports the evidence that this isoform is essential for maintaining spine structure and function [3]. Furthermore, at postsynaptic sites we observed a poor co-localization of the NF isoforms, suggesting that they do not exclusively assemble in hetero-filaments. Together with the demonstration of isoform-specific interaction with receptors at synapses (e.g. NFL and NMDA receptors, NFM and D1 dopamine receptors, [10,12,19]), this data points at isoform-specific functions of NFs within the context of the postsynapses.

To follow up on the hypothesis that different NF isoforms have different functions at the postsynapse, we investigated whether their presence depends on synaptic traits such as synaptic strength or activity. These analyses showed that the postsynaptically less abundant isoforms NFM and NFH also show lower correlation with synaptic strength and activity. The former has been shown to interact with dopaminergic receptors. Since in primary hippocampal cultures the majority of the population is represented by glutamatergic and not dopaminergic neurons [27], the low level of correlation for NFM is unsurprising, and further is in agreement with evidence that NFM-null mice exhibit normal neurotransmission and LTP induction [3]. In NFH-null mice, however, LTP is impaired via yet unknown mechanisms. The lower level of correlation between NFH amount and presynaptic activity and postsynaptic strength could therefore suggest its additional involvement in either activity independent synaptic processes or in more long-term synaptic plasticity mechanisms. Notably, the epitope recognized by the NFH antibody (AA998-1097 of mouse NFH [28]), presents only a single phosphorylation site and therefore it is unlikely that the reduced correlation is due to different posttranslational modifications.

NFL showed a slightly different behavior than NFM or NFH, and did moderately correlate with both, synaptic activity and strength. This indicates that more active synapses had higher amounts of NFL present at the postsynapse, suggesting a contribution of NFL to short-term plasticity and immediate response to activity changes, as well as an involvement in long-term structural responses that would eventually result in an increased synaptic strength. Since NFL is known to interact with NMDA receptors via the GluN1 subunits the correlation between synaptic activity and NFL amount could also demonstrate its direct involvement in the modulation of synaptic signaling [16]. As a cytoskeletal component the involvement of NFL might include NMDA receptor positioning or more indirect changes mediated by the actin cytoskeleton.

Concluding, in this study we systematically quantified the presence of NFs at the postsynapse within the context of the synaptic status. Further work will be needed to clarify the behavior of internexin, the mechanisms through which NFL responds to synaptic activity, and how the presence of NFM and NFH at the postsynapse is regulated.

## Author Contributions

Conceptualization, E.D.; methodology, G-M.G. and E.D.; software, M.A.R.B.F.L., A.R.C.D.; validation, C-M.G., M.A.R.B.F.L., E.D.; formal analysis, C-M.G., M.A.R.B.F.L.; investigation, G-M.G.; resources, A.R.C.D, V.M.P., J.H.; data curation, E.D.; writing—original draft preparation, C-M.G., M.A.R.B.F.L., E.D; writing—review and editing, all authors.; visualization, G-M.G.; supervision, E.D.; project administration, E.D.; funding acquisition, E.D. All authors have read and agreed to the published version of the manuscript.

## Informed Consent Statement

Not applicable.

## Data Availability Statement

Raw imaging data is made available upon reasonable request.

## Acknowledgments

We thank Annette Herold, Magnus-Carsten Huppertz and Birgit Koch for support with neuronal cultures and virus production. We thank Jade Cottam-Jones for critical reading of the manuscript.

