## Supplementary information for "Synaptic status differentially regulates neurofilaments in dendritic spines"

### Supplementary Method - Western Blot

Whole mouse brain lysate or cultured hippocampal primary neurons (DIV18) lysed in 1x Laemmli buffer (Bio-Rad, cat. 161-0747) were loaded in a 4-15% gradient SDS gel (Bio-Rad, cat. 4561086), together with a protein standard (Bio-Rad, cat.161-0394) and subjected to electrophoresis at 50 V for 5 min, followed by 100 V for 90 min. Separated proteins were then wet transferred onto a PVDF membrane (Bio-Rad, cat. 162-0260) at 120 V for 60 min. The blotted membranes were blocked in Tris-buffered saline, 0.1% Tween (TBST), 3% BSA and incubated overnight with previously described NF primary antibodies and for 1 hour at RT with α-tubulin mouse (Sigma, cat. T9026) or GAPDH rabbit (Cell Signaling, cat. 2118) at 1:500-1:1000 dilution in TBST, 1% BSA. Then the membranes were washed five times in TBST and incubated with secondary antibodies (Alexa Fluor 488 anti-mouse, Thermo Fisher, cat. A32723, Alexa Fluor 488 anti-chicken, Thermo Fisher, cat. A-21467, Alexa Fluor 647 anti-rabbit, Thermo Fisher, cat. A-21245) at 1:1000 dilution in TBST 1% BSA for 1 hour at room temperature. Afterwards membranes were washed five times in TBST and fluorescent signal was detected in a ChemStudio (Jena Analytik GmbH).

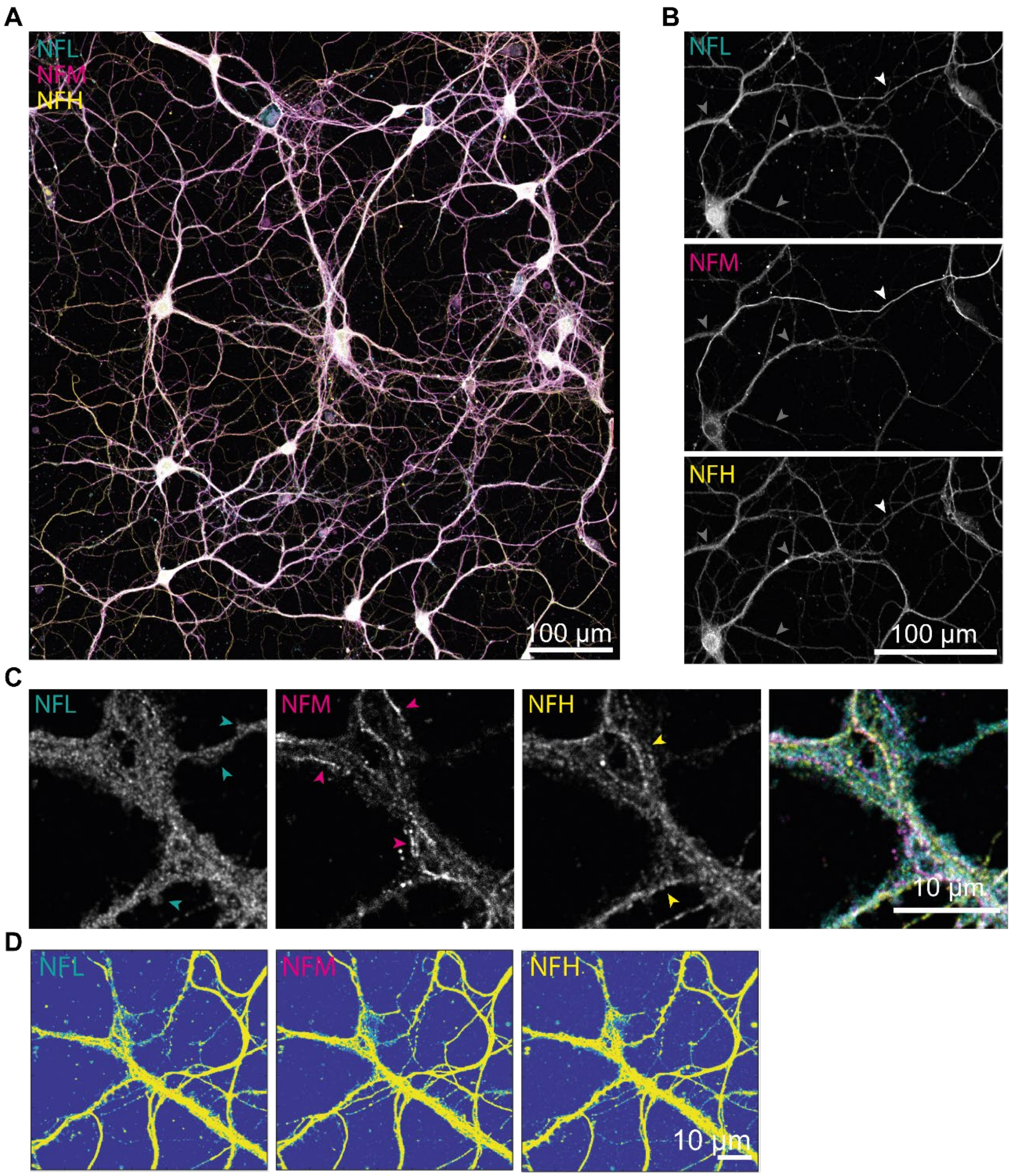


**Figure S1. Immunofluorescent labeling of NFs in primary hippocampal cultures reveals different distribution pattern.** (A) Confocal image of the different NF isoforms NFL (cyan), NFM (magenta) and NFH (yellow) in primary cultured neurons. Scale bar is 100 µm. (B) Enlarged single channel image of panel A. White arrow points at filamentous axonal signal, gray arrows at dendritic regions with more dispersed signal. (C) Enlarged area of Figure 1A. Arrows in single channel images indicate characteristic granular or filamentous NF structure. (D) Representative segmentation by thresholding of individual confocal fluorescent channels for colocalization analysis. Images correspond to the field of view shown in Fig. 1A. Only yellow areas in the cellular regions are considered for the colocalization analysis.

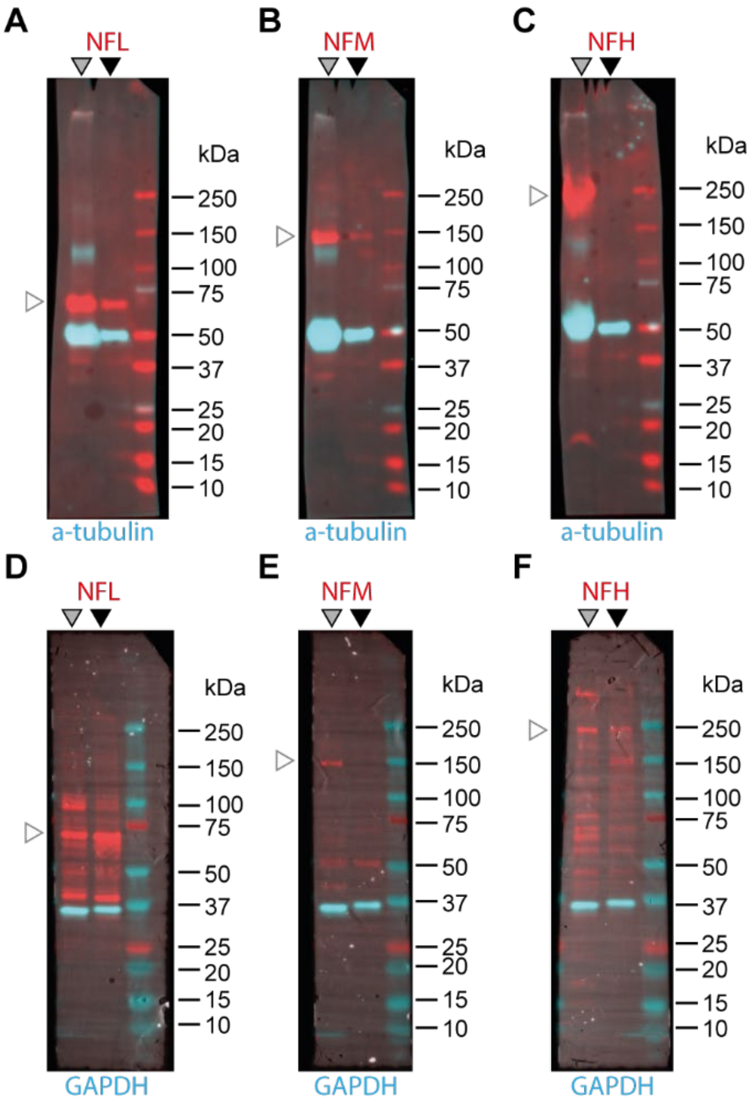


**Figure S2. NF antibody validation in western blot.** Dual color fluorescent western blot on whole brain lysate of a P18 mouse (left lane, gray arrow) or lysate of rat hippocampal cultures as used for STED experiments (middle lane, black arrow), compared to a protein standard ladder (right lane). Top row: NFL (A) NFM (B) or NFH (C) rabbit primary antibody in red. Alpha-tubulin mouse primary antibody in cyan. Bottom row: NFL chicken (D), NFM mouse (E) or NFH mouse (F) primary antibody in red. GAPDH rabbit primary antibody in cyan. White arrows show expected heights of NF isoforms. Please note that according to the manufacturer the NFH antibody used in C is not recommended for WB.

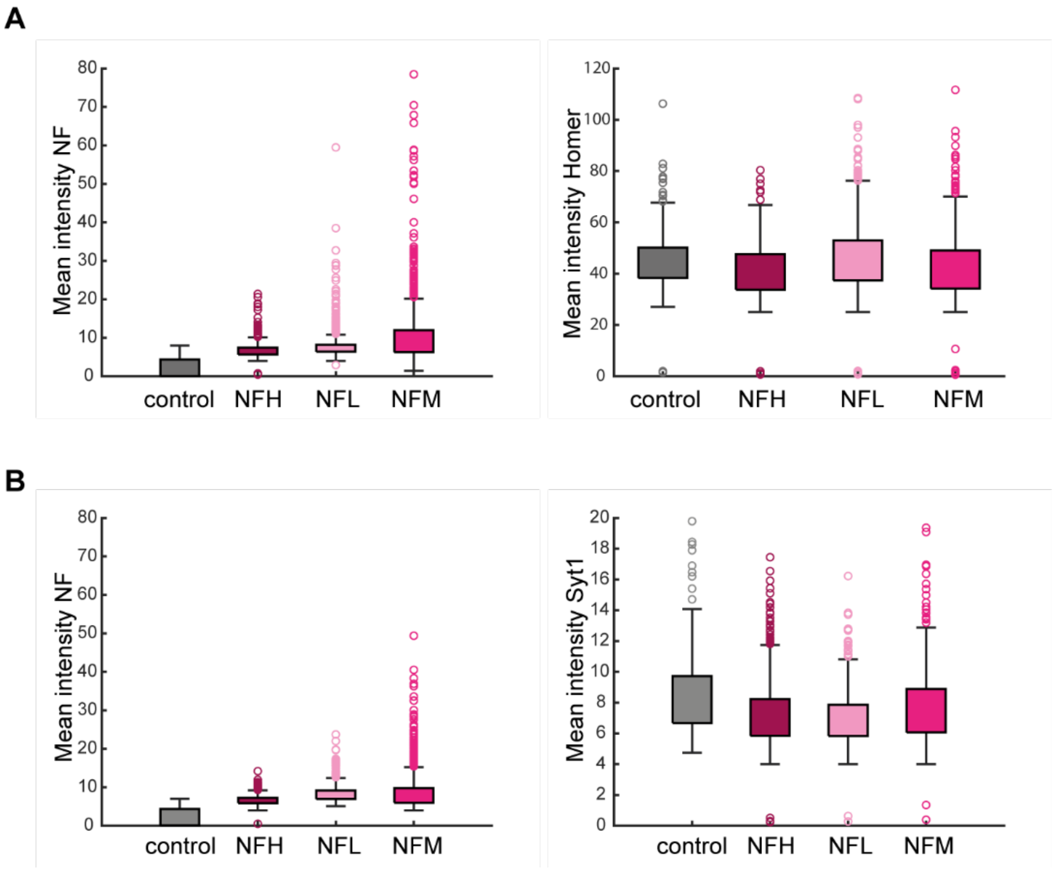


**Figure S3. Intensity quantification of NF and synaptic labeling at manually segmented postsynapses in dual color STED images.** (A) Mean intensity of NF (left) and homer (right) in manually segmented spines from 6 independent experiments. (B) Mean intensity of NF (left) and Syt1 (right) in manually segmented spines from 3 independent experiments. Controls are samples labeled without primary NF antibodies. Values represent mean of raw photon counts.
